# The mechanical basis for snapping of the Venus flytrap, Darwin’s ‘most wonderful plant in the world’

**DOI:** 10.1101/697797

**Authors:** Jan T. Burri, Eashan Saikia, Nino F. Läubli, Hannes Vogler, Falk K. Wittel, Markus Rüggeberg, Hans J. Herrmann, Ingo Burgert, Bradley J. Nelson, Ueli Grossniklaus

**Author notes:** these authors contributed equally to this work (Jan T. Burri, Eashan Saikia, Nino F. Läubli, Hannes Vogler).

## Abstract

The carnivorous Venus flytrap catches prey by an ingenious snapping mechanism. Based on work over the past 190 years, it has become generally accepted that two touches of the trap’s sensory hairs within 30 seconds, each one generating an action potential, are required to trigger closure of the trap. We developed an electromechanical model which, however, suggests that under certain circumstances one touch is sufficient to generate two action potentials. Using a force-sensing microrobotics system, we precisely quantified the sensory hair deflection parameters necessary to trigger trap closure, and correlated them with the elicited action potentials *in vivo*. Our results confirm the model’s predictions, suggesting that the Venus flytrap may be adapted to a wider range of prey movement than previously assumed.

## Introduction

The hunting mechanism of the carnivorous Venus flytrap (*Dionaea muscipula*), according to Darwin ‘the most wonderful plant in the world’^1^, has attracted the interest of many scientists, starting with the observations made by Moses A. Curtis in the 1830s.^2^ Since then, the individual phases from trap triggering to the reopening after successful digestion have been investigated from different angles (reviewed in ^3^). Starving plants attract insects through the secretion of volatile compounds.^4^ While exploring the trap for food, wandering insects accidentally touch one of the six sensory hairs distributed on the two lobes of the trap, thus triggering an action potential (AP).^5–7^ A second touch-triggered AP within about 30 seconds causes the trap to snap, and the prey is caught. Further APs triggered by the struggling prey induce jasmonic acid biosynthesis and signalling,^8, 9^ which seals the trap tightly and eventually leads to the formation of the ‘green stomach’, a digestive cocktail that mobilizes prey-derived nutrients.^7, 10, 11^

In this paper, we focus on the translation of the mechanical stimulation of the sensory hairs into an electrical signal. Although there is a general agreement that the sensory hair deflection opens mechanosensitive ion channels, such channels have not been identified yet.^12, 13^ While these putative channels are open, a receptor potential (RP) builds up.^6, 14^ If the deflection of the sensory hair is large enough, the RP reaches a threshold, above which an AP is generated. Jacobson tested the generation of RPs and APs with respect to mechanical stimuli of different magnitudes in an *in vitro* system using dissected traps, and observed a correlation between the rate of the stimulus and the generation of APs.^6^ However, the experimental setup available at the time did not allow an exact quantification of the deflection parameters nor the observation of trap closure. We used a microelectromechanical (MEMS)-based force sensor mounted on a microrobotics system to precisely control the velocity and amplitude of the deflection and to simultaneously measure the applied force. In this way, we were able to accurately quantify the parameter range in which hair deflection leads to trap closure, while a second force sensor measured the generated snapping force. In addition, we measured APs, using a non-invasive method, to test the deflection conditions under which they are generated. An electromechanical model inspired by the literature^15^ was fitted with our own data and suggested that, in the case of slow mechanical stimulation, a single deflection is sufficient to generate multiple APs. We experimentally confirmed this prediction and show that, contrary to the general assumption, a single, slow hair deflection leads to the generation of two APs that trigger trap closure.

## Methods and Materials

### Venus flytraps

Our Venus flytrap population of about 100 plants was originally grown from seeds in 2011. The original seed batch was a donation from the Botanical Garden Zurich (https://www.bg.uzh.ch/de.html). Once a year the plants were split and re-potted into 9 cm clay pots, which were reused after they had been cleaned from moss. As a substrate, we use a mix of 90% white peat (Zürcher Blumenbörse, Wangen, Switzerland) and 10% granulated clay (SERAMIS Pflanz-Granulat, Westland Schweiz GmbH, Dielsdorf, Switzerland). The Venus flytraps were grown in a greenhouse with 60% relative humidity and a temperature regime of 18 to 23°C during the day and 16 to 21°C during the night. Plants were grown under normal daylight with morning and evening periods extended by 400 W metal-halide lamps (PF400-S-h, Hugentobler Spezialleuchten AG, Weinfelden, Switzerland) to ensure a daylength of 16 h. The lamps were also turned on when the daylight was not sufficient. Rain water was used to irrigate the plants whenever possible. For the experiments, the plants were transported in an insulated moisture chamber from the greenhouse to the laboratory. All experiments were performed *in vivo*. Plants were put back in the moisture chamber after every set of measurements.

### System configuration

The system consists of two subsystems, the first of which combines a force probe with a microrobotics system to quantify the sensory hair deflection parameters. The second subsystem uses a load cell, metal levers, and hinges to measure the snapping forces of the trap (Fig. 1a). The second subsystem also served to protect the force probe of the first subsystem from damage by preventing the trap from closing when the snapping mechanism was triggered. In addition, we attached the electrode for measuring the APs to the metal levers, such that it was unnecessary to re-attach the electrode to individual traps.

**Figure 1.**
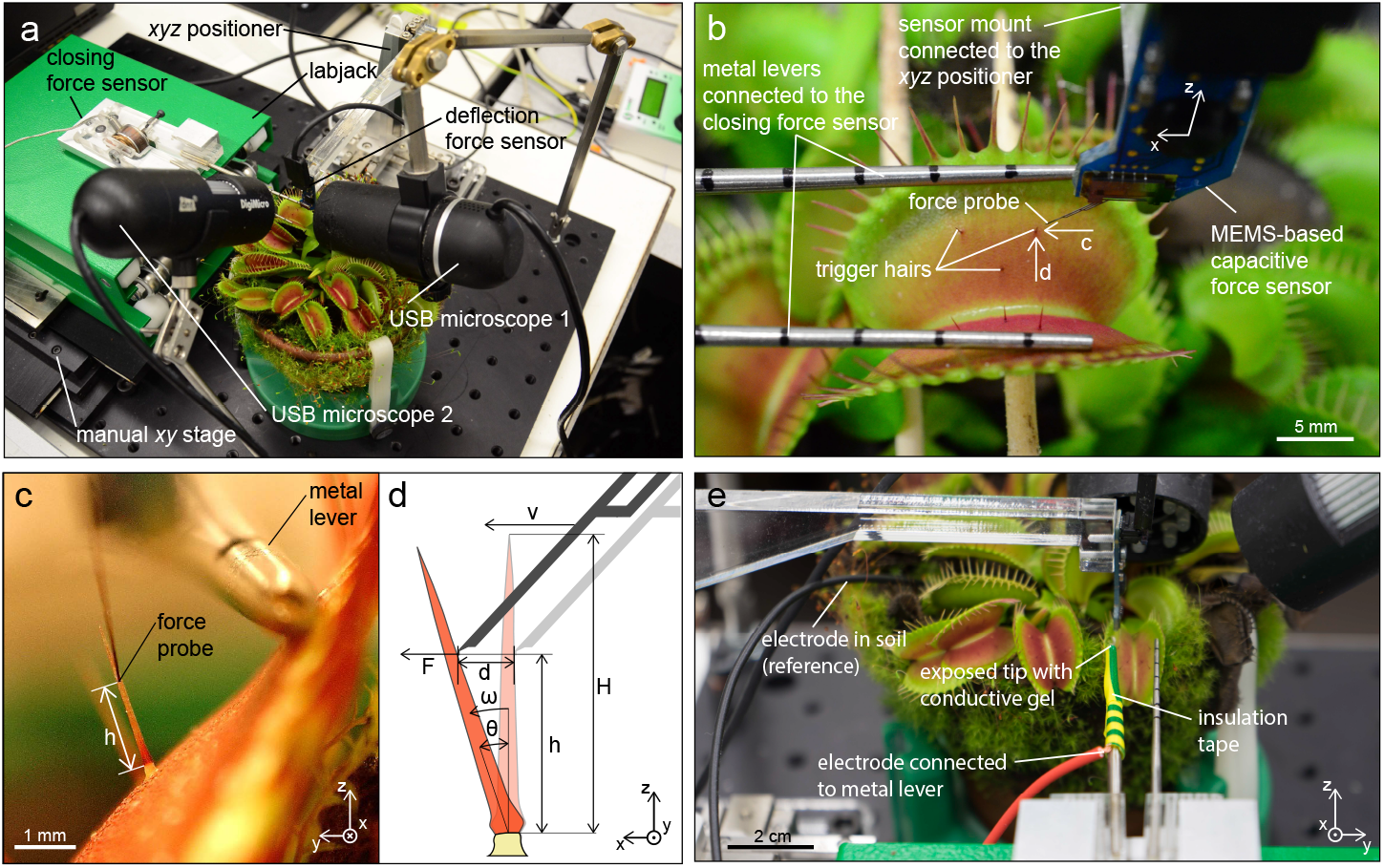
System configuration to investigate the necessary mechanical stimulation of the sensory hair to initiate trap closure while measuring the resulting forces and action potentials (APs). a) The system combines a force probe to precisely control the deflection of sensory hairs and a second force sensor that measures the subsequent closing forces. Two USB microscopes are used to observe the experiment. (b) Close up of the deflection force sensor used to deflect the hair in direction parallel to the midrib. (c) View from the USB microscope 1 the right in (a) used to extract the geometry of the sensory hair. (d) Schematic side view of the hair deflection where the velocity *v*, the deflection *d* and the force *F* is controlled and monitored. (e) For the AP measurements one electrode was connected to the metal lever and the other one inserted into the soil.

### Sensory hair deflection

The deflection force sensor is a MEMS-based capacative force sensor (FT-S1000-LAT; FemtoTools AG, Buchs, Switzerland) with a force range of ±1000 μN with a standard deviation of 0.09 μN at 200 Hz, which measures the forces applied to a force probe (50×50 μm) in *x* direction. The force signal was recorded with a multifunction I/O device (NI USB-6003; National Instruments (NI), Austin, TX, USA). The deflection force sensor was mounted via a custom made acrylic arm to a *xyz* positioner (SLC-2475-S; SmarAct, Oldenburg, Germany) with a closed-loop resolution of 50 nm.

The deflection force sensor was placed inside the trap with the force probe in front of a sensory hair guided by the optical feedback of two USB-microscope cameras (Fig. 1a, b). The view from the USB-microscope 1 (DigiMicro Profi; DNT, Dietzenbach, Germany) is used to position the probe laterally in the centre of the hair and to extract the length of the sensory hair *H* and the distance *h* of the contact point of the sensor probe relative to the constriction (Fig. 1c). For that purpose, a reference step of 500 μm was performed with the force probe in *z* direction, and the pictures captured at start- and endposition were used to determine the distance-per-pixel value. The USB microscope 2 (DigiMicro; DNT), oriented perpendicularly to the first one, was used to bring the force probe in close proximity to the sensory hair. Additionally, the force sensor could be automatically placed at a defined distance from the hair by finding contact using the force readout and then moving back to a defined position.

We defined a single deflection as a combination of back and forth angular displacement of the hair caused by the advance and return of the force probe in *x* direction (i.e. parallel to the midrib). This movement is parallel to the force-sensitive direction of the force probe and also ensures a perpendicular contact between force probe and sensory hair. The velocity of the sensor probe *v* during deflection can be defined and either a maximum force *F* or a maximum linear displacement *d* is set to define the maximum deflection (Fig. 1d). The initial angular velocity *ω* is then given as

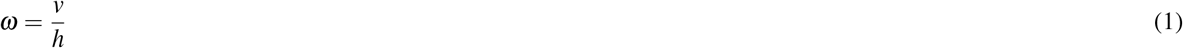

Furthermore, the angular displacement *θ* during the deflection is given by

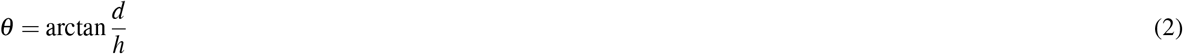

The applied torque *τ* is given by

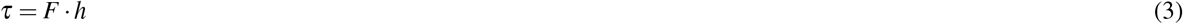

Feedback control and data logging of forces and positions during the deflection were executed in LabVIEW™.

### Action potential measurement

The action potentials were recorded by connecting an electrode to one of the metal lever arms of the closing force sensor and inserting the reference electrode into the soil (Fig. 1e). Insulating tape is used to cover most of the lever only exposing the tip which is in contact with the trap leaf. A droplet of conductive gel (Elektroden Gel 250G, Compex) was applied between tip and leaf to guarantee a better contact area stabilizing the readout and increasing the signal. The voltage between the two electrodes was read out using an analog input module (NI USB-6003; NI).

### Closing force measurements

The closing force sensor consisted of two metallic lever arms, which transferred the force of the closing trap onto a load cell (31E Mid; Althen Sensors & Controls, Leidschendam, Netherlands) with a load capacity of 50 N and a resolution of 75 mN (Fig. 2a). Both arms were hinged at one end, thus opening and closing similar to a second order lever. The load cell was fixed onto one of the lever arms, while an adjustable screw was inserted into the other lever arm at the same distance *l*_*cell*_ from the hinge. The gap between the two levers could then be adjusted by loosening or tightening the screw in order to measure flytraps of variable sizes and shapes. The closing force sensor was mounted on a labjack and placed on a manual *xy* stage. This setup allowed the precise spatial positioning of the lever arms within the trap.

**Figure 2.**
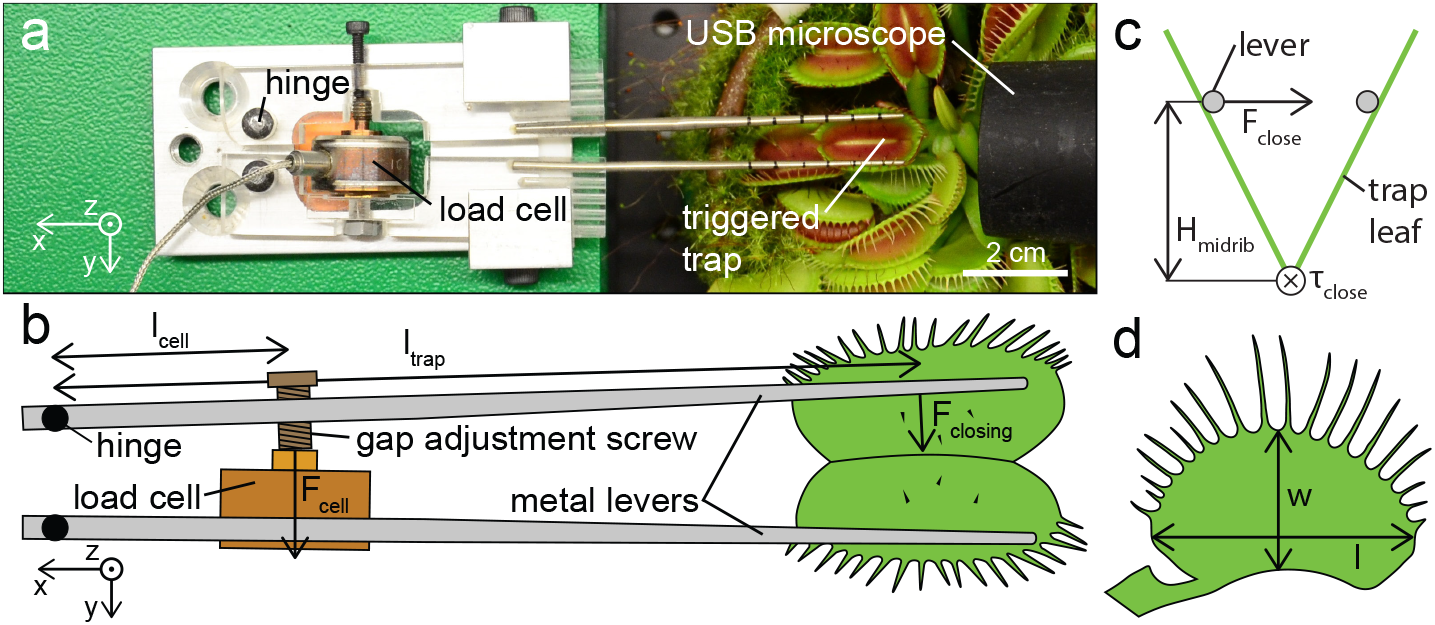
Top view of the closing force sensor with a triggered trap. (b) Schematic top view of the closing force sensor showing how the force generated by the flytrap is transmitted onto the load cell. (c) Relation between measured force *F*_*close*_, the height of the levers and the closing torque *τ*_*close*_. (d) Schematic of the measured geometries, i.e. width *w* and length *l* of a trap.

The relation between the force *F*_*close*_ applied at the lever by a closing trap and the resulting force at the load cell *F*_*cell*_ (Fig. 2b) is given by

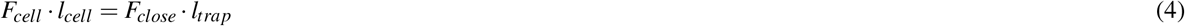

where *l*_*cell*_ and *l*_*trap*_ are the distances from the hinge to the load cell and to the middle of the trap leaf, respectively. The closing torque *τ*_*close*_ is given by

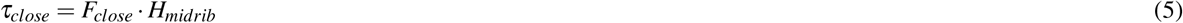

where *H*_*midrib*_ is the height between midrib and metal lever (Fig. 2c). *H*_*midrib*_ was determined using the ratios between width and length of the trap leaf (Fig. 2d) and the top-down images of every individual experiment. The width and the length of 31 leaves were measured with a caliper to determine the ratio. The ratio *w*/*l* of the leaves had a mean value of 0.59 and a standard deviation of 0.04. The mean value was used to calculate the torque for the remaining 16 experiments where we did not measure the geometries.

The lever arms were carefully moved into the trap of a Venus flytrap, while maintaining an optimal gap between them, in order to prevent contact with the sensory hairs. The gap between the levers was then increased until they contacted the surfaces of the lobes on both sides and underneath the marginal teeth. This configuration was maintained throughout the sensory hair deflection experiments where it reduced perturbations by fixing and stabilizing the two lobes of the trap. By keeping the two lobes of the trap open even after closure had occurred, we could ensure that the deflection force sensor would not be damaged. The force signal was continuously recorded using a analog input module (NI USB-6009; National Instruments (NI), Austin, TX, USA).

Additionally, the time between mechanical stimulation and trap snapping (i.e. delay time) was determined by counting the frames recorded by the USB microscope between the second deflection of a hair and the first movement of the trap.

### Empirical charge-flow (ECF) model

It has been previously observed that the trap tissue is capable of accumulating subthreshold charges, and different types of “electrical memory” were identified, such as the sensory memory, short term memory and long term memory, by external injection of charges with a capacitor.^16^ If the total charge transmitted by the capacitor exceeded 14 μC, the trap was found to close.

Based on these descriptions, we introduce an empirical model to gain a deeper understanding for the charge-flow and its accumulation behaviour within the Venus flytrap. The objective of this model is to qualitatively reflect on the set of *in vivo* experimental observations by considering the simplest version of biologically equivalent resistor-capacitor (RC) circuit for the Venus flytrap, as described in^15^. The two types of electrical memory that are relevant to our model, are the sensory memory and short-term memory. This is because, our model captures the effects that occur only during the time scale of a stimulus, whereas the long term memory corresponds to the information storage throughout its entire lifetime. The sensory memory immediately follows a stimulus and then, the information is relayed to the short term memory via the flow of charges. The stimulus parameters of our deflection experiments, namely the angular velocity *ω* and the angular deflection *θ*, act as the input parameters for our model, while our analytical relations are based on the sensory memory and the short term memory, both defined in terms of decay time constants *τ*_*sm*_ and *τ*_*stm*_ respectively. The decay of charge in the sensory memory and short term memory is modeled as an exponential decay, as in the case of a simple RC circuit. We consider that both the advance and return phases of the hair bending during a single deflection, has the same effect on the charge addition and accumulation. This is because, the plant does not differentiate between advance and return of the probe, which was made clear by the fact that electrical pulses were detected at different instances of the hair bending in different plants. Hence, the total time of excitation *T* in our model, comprises of both the advance and return phases of the force probe, and it is defined as *T* = 2*θ/ω*.

*T* is then discretised into a finite number of time steps *n*, each with a fixed step size of Δ*t* seconds, such that *n* = *T/*Δ*t*. During every time step, a fixed amount of charge *Q* is freshly added to the sensory memory which is expressed as a function of *ω*, such that *Q* = *f* (*ω*). Our hypothesis behind this expression is that, a hair would undergo a larger deflection at higher angular velocities during a fixed time step. Consequently, a larger deformation is produced on the cell walls and membranes of the sensory cells, which leads to the activation of more mechanosensitive ion channels. As fresh charge is added, the total charge in the sensory memory at a particular time step can be expressed as the summation of the freshly added charge *Q* and the remaining charge from the previous time step, after its decay.

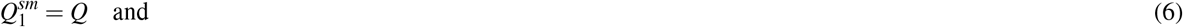

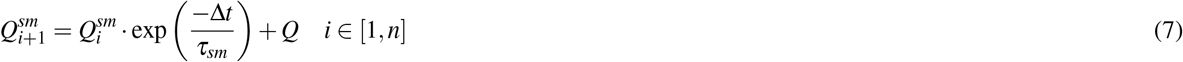

A fraction *β* of this charge content in the sensory memory is then transferred into the short term memory.

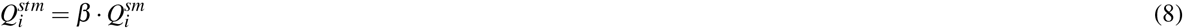

and the cumulative charge during a time step can be expressed as

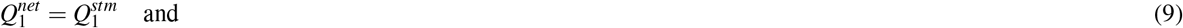

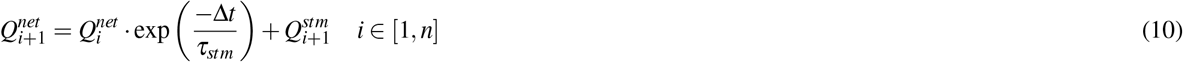

In the event that a second deflection is imparted as a successive input stimulus, the charge decay in between the two stimuli is influenced by the memristive behaviour of the leaf tissue during the time *t*_*gap*_ in between two successive deflections. This is incorporated into this model by defining a typical non-exponential decay function *R*_*p*_ as observed in the case of ideal memristive-capacitive circuits.^17^

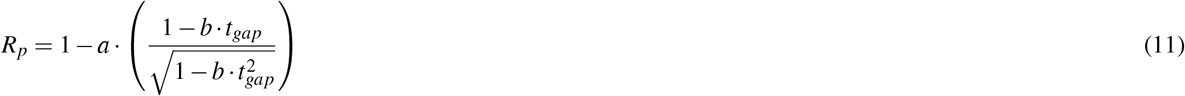

Following this, the second stimulus has the same contribution as the first stimulus, where the additional charge is accumulated in the sensory memory and the short term memory. The model was formulated in MATLAB 2018b and it was evaluated for different combination of stimulus parameters, *ω* ∈ [0.001,10] rad/s and *θ* ∈ [0,1] rad.

## Results

### Two fast consecutive deflections trigger trap snapping if a certain angular displacement or torque is reached

Prey animals wandering on the trap trigger the snapping mechanism by touching the sensory hair(s) twice within 30 s. To simulate these stimuli, we deflected the hair twice at maximal positioner speed, which resulted in high initial angular velocities, typically ranging from 10 to 20 rads^−1^ (depending on the contact height *h* of the force sensor probe on the sensory hair). During the deflection, we quantified the forces necessary to sufficiently bend the sensory hair to activate the trap. We defined a single deflection as the combination of a back and forth angular displacement, similar to what happens when an animal touches the hair. The measurements were performed on 21 sensory hairs, each on a different trap from 12 individual plants. Each measurement consisted of two subsequent deflections with a 1 s gap between them, up to a predefined angular displacement *θ*. If the trap did not shut, we waited for 2 min to make sure that trap memory was completely reset, then the experiment was repeated on the same sensory hair but with a larger angular deflection. The waiting time was chosen long enough to allow the dissipation of any possible charge that may have built up due to the previous double deflection experiment. The measurement was repeated with increasing angular deflection until snapping of the trap was triggered (Fig. 3a,b), which was the case when an median angular displacement threshold of 0.18 rad or a median torque threshold of 0.8 μNm (Fig. 3c and Fig. S1a) was reached. Therefore, an animal touching the hair needs to apply around 0.5 mN close to the end (i.e. three quarters from the constriction) of the sensory hair, or up to 5 mN close to the constriction (Fig. S1b).

**Figure 3.**
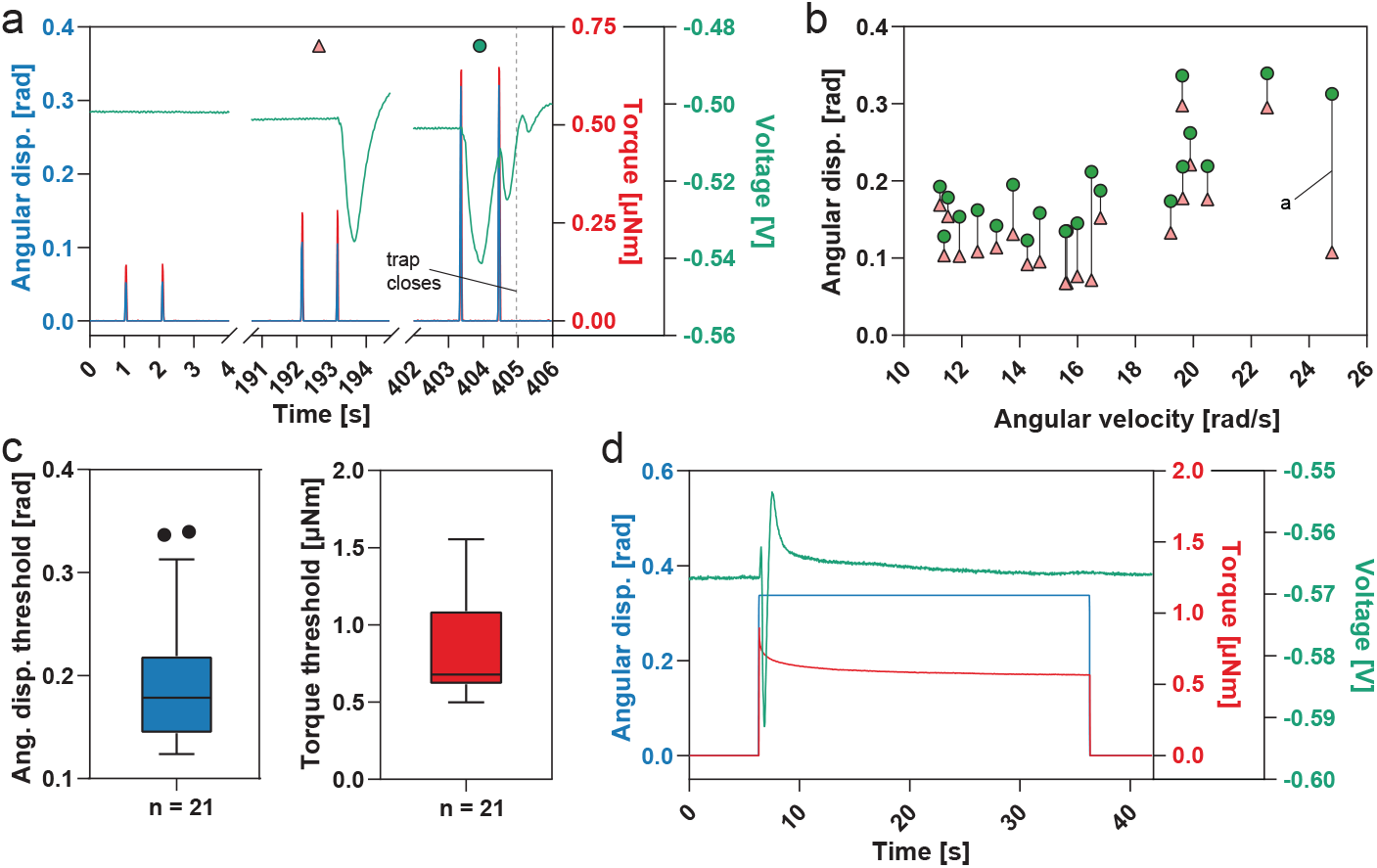
Double deflection experiments and sustained displacements of sensory hairs. (a) Subsequent double deflections of the sensory hair with increasing angular displacement controlled by a sensor probe while the torque and the voltage readout between leaf and soil are recorded. The green circle marks the stimulation that led to trap closure with every deflection eliciting an action potential (AP). The red triangle marks the preceding insufficient stimulation, where the two deflections add up to exceed the receptor potential (RP) threshold leading to a single AP. (b) 17 individual measurements showing the double deflection that led to trap triggering (green circle) and the preceding double deflection that did not reach the threshold (red triangle). The experiment shown in (a) is indicated. (c) Descriptive statistics of the angular displacement threshold and torque threshold for double deflections that initiated trap closure (i.e. resulted in two AP being fired). The middle line indicates the median, the box the 25^*th*^ and 75^*th*^ percentile and the whiskers extend to the most extreme data points not considered outliers. (d) A 30 s sustained angular displacement of the sensory hair with the recorded torques and the voltage readout on the leaf. The initial deflection leads to an AP, but the sustained displacement did not contribute to the RP.

AP measurements provide the missing link between hair deflection and trap closure. Whether an AP is generated depends on the RP, which has to reach a critical threshold.^6^ When the two consecutive deflections were well below the angular deflection threshold, we never observed an AP. For deflection amplitudes near the angular deflection threshold, a single AP was elicited after the second deflection. This indicates that both deflections contribute to the RP and that the threshold to fire an AP was only reached with the second deflection. As expected, a single AP was not sufficient to trigger trap closure. If the deflections exceed the deflection threshold, two APs, one for each deflection, were generated leading to trap closure. These results suggest that a fast deflection of the sensory hair increases the RP to a certain level, which depends on the amplitude of the angular deflection. RPs can add up and may elicit an AP after several deflections if they are below the deflection threshold. However, the generation of one AP per touch only holds true if sensory hair deflection is above the deflection threshold.

### Sustained angular displacement does not trigger trap closure

Since the increase of the RP from multiple deflections is additive, the question arose whether a sustained displacement had a similar effect. To test this, we deflected 11 sensory hairs, each from a different plant, far beyond the angular displacement threshold, and kept that position for 30 s (Fig. 3d). None of the traps closed during the sustained deflection (mean value *θ* = 0.31). AP measurements revealed that the initial displacement elicited a single AP, after which the voltage quickly returned to the baseline despite the hair staying deflected. If the sustained displacement had contributed to the RP, it would have remained above the threshold, in which case we would have expected a series of APs. To verify that the tested sensory hairs were responsive, we subsequently performed double deflection tests, which triggered trap closure in all cases.

It appears that triggering trap closure does not depend on the displacement of the sensory hair itself, but rather on the rate of the displacement. This is in contrast to a previous finding where prolonged pressing led to trap closure, but manual stimulation.^15^

### An empirical charge-flow (ECF) model suggests that a single deflection is sufficient to trigger trap closure

After quantifying the deflection parameters in the classical situation with two touches, we were wondering about the limits of angular displacement and velocity within which the traps would react. Therefore, we developed an ECF model and evaluated it for the condition that trap closure is triggered as soon as a threshold value *Q_th_* of cumulative charge is exceeded. We chose a threshold *Q_th_* of 8 μC because it was previously reported that an equivalent amount was necessary for smaller plants to initiate trap closure^15^. We first tested the model with single deflections and, to our big surprise, it predicted that a single touch would generate multiple APs under certain circumstances. This suggested that within given boundaries, a single deflection would be sufficient to trigger snapping of the trap.

The regime for all possible combinations of *ω* and *θ* for which the model predicted trap closure by a single touch is marked by small blue dots (Fig. 4). Their respective threshold values lie on the boundary of this region. In a very low angular velocity range (*ω* < 0.01 rad/s), the traps will not shut according to the model. Increasing the value of *ω* eventually led to trap closure, whereby the threshold of angular deflection *θ* that is necessary to initiate trap closure attained a minimum value of 0.05 rad within a range of the angular velocity *ω* between 0.02 and 0.05 rad/s. By further increasing the angular velocity (*ω* > 0.05 rad/s, the model predicted a monotonous increase of the *θ* threshold value required for trap closure until an angular velocity of *ω* > 9 rad/s was reached, after which the trap would not close anymore. Beyond this upper limit, a single deflection is predicted incapable of triggering trap closure, irrespective of the extent of the deflection. As a consequence, a second deflection is then necessary to supplement the charge accumulation from the first deflection stimulus.

**Figure 4.**
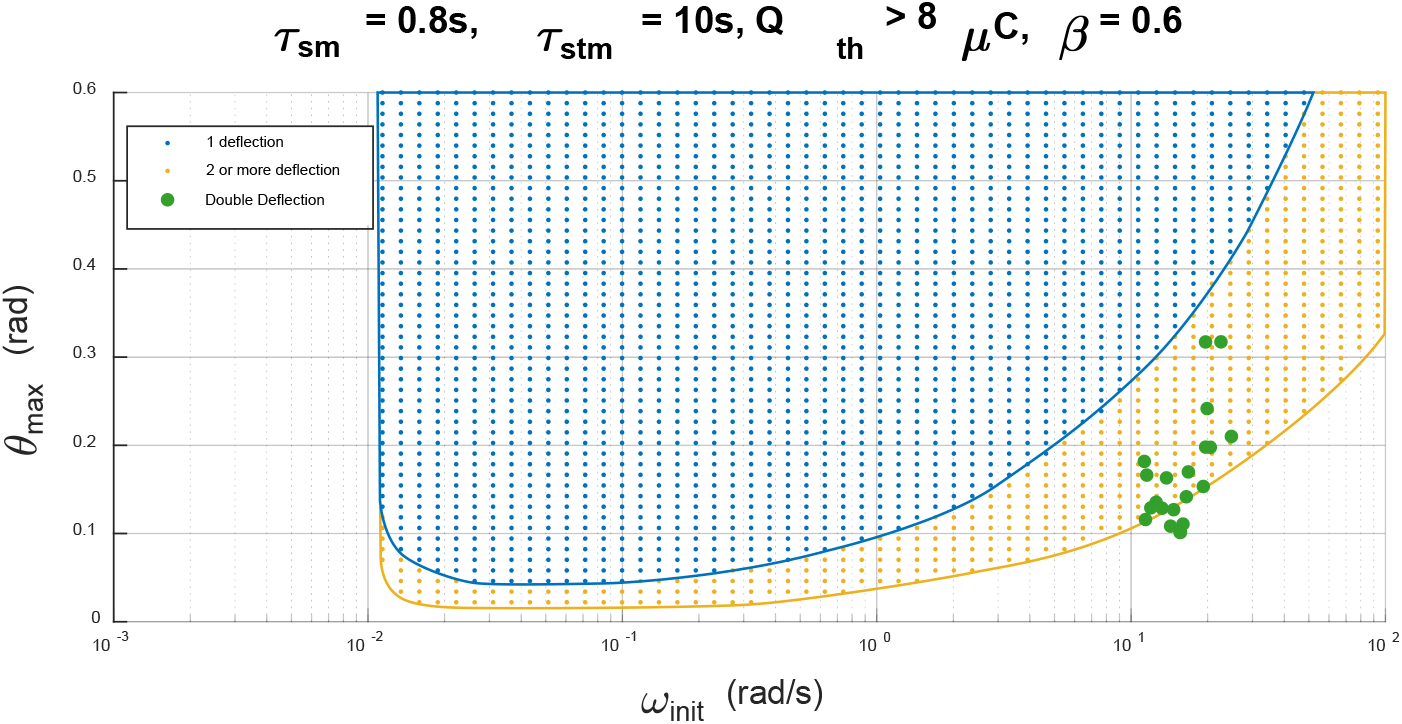
A snapshot of the ECF model output for a set of parameters comprising of the decay constants *τ*_*sm*_ and *τ*_*stm*_ corresponding to the plant sensory memory and short term memory. A fraction *β* = 0.3 of charge is transferred from the sensory memory into the short term memory. Charge threshold *Q*_*th*_ of 8 *μ*C was pre-set as the condition for successful trap closure. Specific zones of successful trap closure by a single deflection (blue dots), two or more deflections (orange dots), separated by solid lines were predicted. Experimental double deflection results are marked by green dots.

### A single deflection at intermediate angular velocity triggers trap closure

Since our ECF model, in contrast to the accepted view, suggested that under certain circumstances a single deflection of the sensory hair would be sufficient to trigger trap snapping, we wanted to test this prediction experimentally. Therefore, we varied the angular velocity of the deflection by adjusting the speed of the sensor probe. Indeed, we were able to confirm the model’s prediction and observed trap closure induced by single deflections at lower angular velocities. To narrow down the range in which this happens, we deflected the same sensory hair several times with varying angular velocities until the trap shut. Between two consecutive deflections, we waited 2 min for the trap to recover and any RP to dissipate. *θ* was chosen as large as possible, up to the point where either slippage between the hair and the force probe occurred or the force range of the force sensor (1000 μN) was exceeded.

The lower boundary of this phenomenon was determined by incrementally increasing the angular velocity *ω* after every deflection (n = 17). A slow initial velocity below 0.009 rad/s was chosen, which never resulted in trap closure. Subsequent single deflections were performed with incrementally increasing velocities until the trap shut (Fig. 5a). The higher boundary was similarly determined by starting with a velocity > 3 rad/s, followed by a step-wise decrease until the trap shut (n = 9). Additionally, we performed another set of single deflection experiments (n = 5) where the angular velocity of the force probe was kept constant at an intermediate speed between 0.2 and 0.4 rad/s, while the angular displacement was gradually increased during subsequent deflections, in order to get the lower boundary of the angular displacement needed for triggering trap closure by a single deflection.

**Figure 5.**
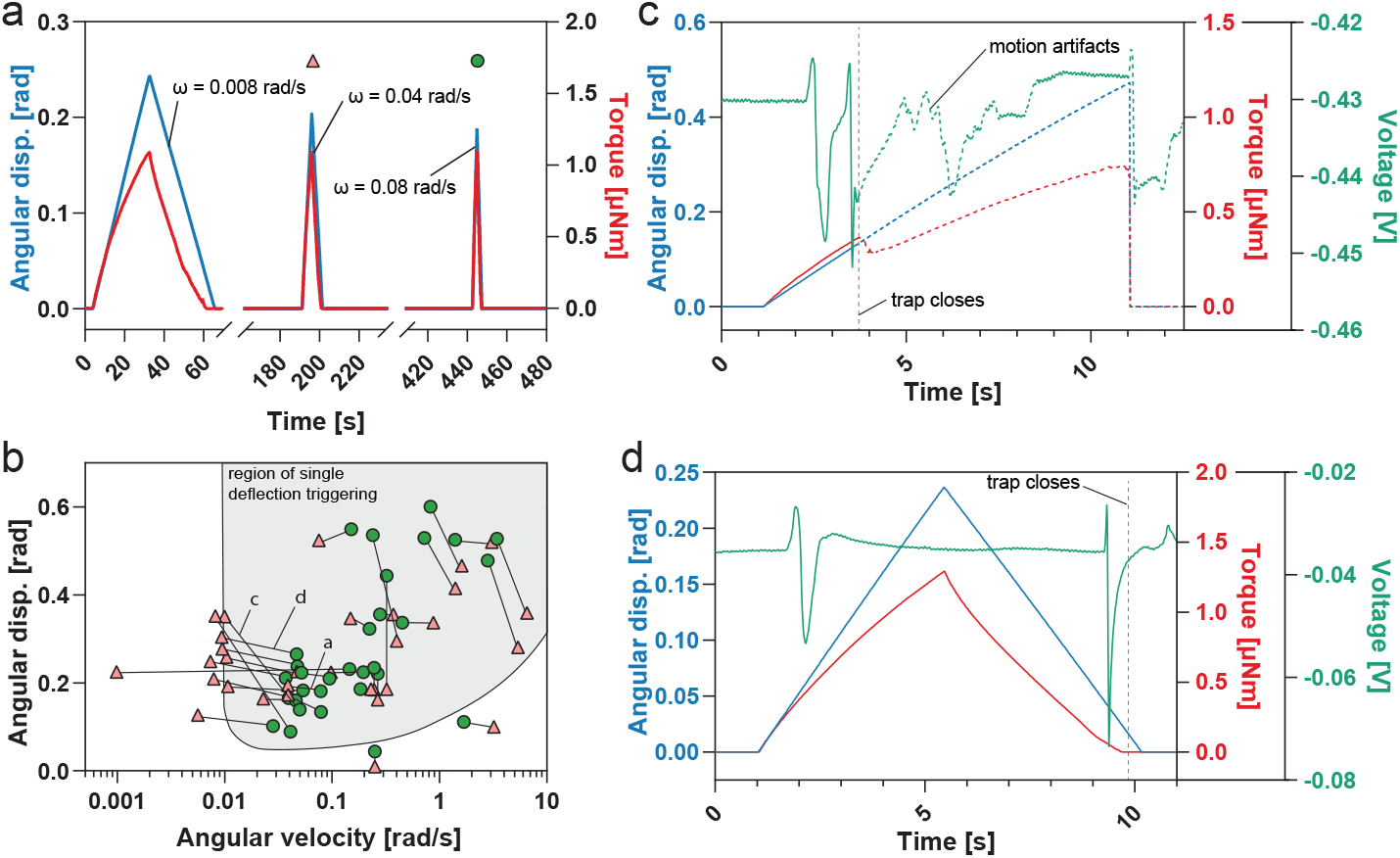
Single deflections at a certain angular velocity lead to trap closure. (a) Subsequent single back and forth displacements of the sensory hair with increasing linear velocity controlled by the sensor probe while the torque is recorded. The green circle marks the stimulation that led to trap closing and the red triangle the preceding, non-sufficient single deflection. (b) Single deflection experiments on 31 individual sensory hair with either increasing or decreasing velocities, or increasing deflections with the stimulus that initiated trap closure (green circle) and the preceding, not successful deflection (red triangle). The experiments from (a), (c) and (d) are indicated. The gray area represents the model output for single deflection triggers. (c,d) Single deflections at an intermediate velocity that led to trap triggering where either (c) two action potentials (APs) are emitted during bending (on the way in), or (d) one AP is released at the beginning of the bending (on the way in) and a second one at the end of the return (on the way out). Trap closure led to motion artifacts in the voltage readout and the resulting change of the hair’s position can affect the determined angular displacement and torque values (indicated by dotted lines).

All the single deflections that resulted in trap closure, together with the preceding stimuli for which no trap closure was observed, define the region of single deflection triggering (Fig. 5b and Fig. S1c). The output from the model for single deflection triggers (marked as gray area) covers a similar parameter space obtained for the angular deflection vs. angular velocity. We observed that a single deflection can be sufficient to trigger trap closure at an intermediate angular velocity of the deflection (0.03 rad/s ≲ *ω* ≲ 4 rad/s), but not sufficient at slower or faster angular velocities. The different angular velocity regimes for our single and double deflection experiments are shown in figure S2.

In the 17 slow-velocity deflection experiments we were able to determine whether trap triggering happened while advancing the force probe (n = 6) or during retraction (n = 11). Our results indicate that both back and forward displacement add up to the overall RP level. AP measurements disclosed how these single deflections lead to trap closure. When the trap closed during the advancement of the force probe, two APs were observed one shortly after the other during the bending of the hair (on the way in) (Fig. 5c). In the second case (triggering during retraction), one AP was fired during initial bending and a second one during retraction of the sensor probe when the hair returned to its original position (Fig. 5d). In both cases, the second AP led to the immediate closure of the trap.

### Quantification of the trap snapping forces and torques

The closing forces were measured at the edge of the lobes where the marginal teeth extend, after the trap was triggered by double deflection (Fig. 6a), and in the single deflection experiments. A median closing force *F*_*close*_ of 73 mN was determined from 48 different traps (Fig. 6b). However, a closing torque *τ*_*close*_ around the midrib is a better quantity to characterize trap closure, since the measured force depends strongly on the position of force readout, the orientation of the leaf, and the size of the leaf. The obtained median *τ*_*close*_ was 0.65 mNm (Fig. 6c). The delay time of the Venus flytrap, i.e. the time between mechanical stimuli and the start of trap closure, was evaluated in double deflection experiments. A mean value of 0.6 s with a standard deviation of 0.3 s (n = 18) was observed, which is in the same range as the 0.4 s previously reported.^18^

**Figure 6.**
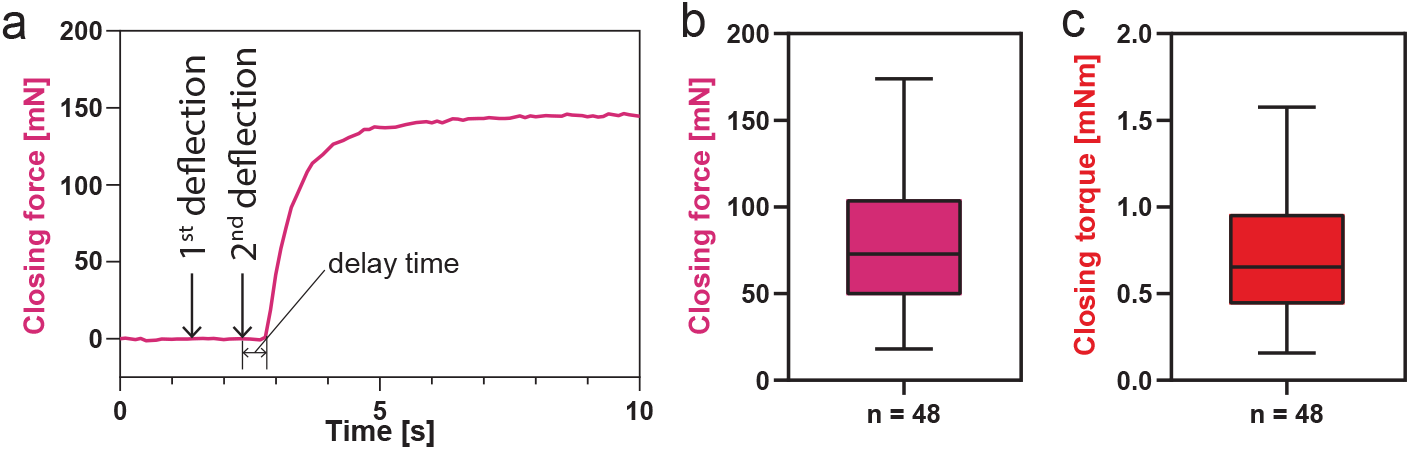
Closing force and torque measured with the closing force sensor. (a) Typical force curve after trap closure induced by double deflection. (b) Descriptive statistics of the closing force right under the marginal teeth and (e) the closing torque around the midrib of a triggered flytrap. The middle line indicates the median, the box indicates the 25^*th*^ and 75^*th*^ percentile and the whiskers extend to the most extreme data points not considered outliers.

## Discussion

We provide novel insights into the first steps of the hunting mechanism of the Venus flytrap, the translation of the mechanical into an electrical signal that eventually leads to trap closure. The generally accepted view is that two touches of the sensory hair by prey animals are necessary to trigger the snapping mechanism. A single deflection of the sensory hair opens putative mechanosensitive channels thereby building up an RP. If the deflection is strong enough, i.e. if a wandering insect touches the sensory hair, the RP reaches a threshold necessary to generate an AP. This causes a transient, subthreshold rise in cytosolic Ca^2+^ levels^3, 7^, which will only surpass the threshold and induce snapping if a second AP follows within about 30 s of the first. Most studies of the correlation between the mechanical stimulus and the electric signalling cascade to date were based on manual deflection of the sensory hair and thus non-quantitative. Semi-quantitative approaches were limited by the technology at the time.^6^ For the first time, we provide a precise quantification of the deflection parameters, such as angular displacement and velocity, which are necessary to trigger the snapping mechanism. In addition, we used a load cell to measure the force that is exerted by the trap upon closure, which is 73 mN but ranges from 18 mN to 174 mN in extreme cases. Therefore, we could largely confirm similar experiments where a piezoelectric sensor film was used instead of a load cell, and which provided average values for the closing force between 140 and 150 mN.^19^ However, the force was measured at the end of the snapping whereas we measured it at the beginning, which is the force needed to restrain an insect until the trap is fully closed.

Reasoning that hair deflection in nature, induced by the legs of spiders, ants and flies, i.e. the ‘classical’ prey of *Dionaea muscipula*,^20^ would be rather quick and snappy, we operated the microrobotics system at full speed in a first experiment. Our observations showed that a minimal angular deflection of 0.1 rad was necessary to trigger an AP for fast deflections with angular velocities between 10 and 30 rad/s. Inspired by the work of Volkov^15^ we developed a simple, electromechanical model and fitted it with our own data. Shifting the deflection parameters beyond the limits of our experimental data, the model suggested that at a lower angular velocity, a single sensory hair deflection should be sufficient to trigger several APs. This is in contrast to the generally accepted idea that every sensory hair deflection generates only one AP. Nonetheless, when performing the corresponding experiments *in vivo*, we indeed found an angular velocity range between 0.03 and 4 rad/s, where a single deflection generated two APs triggering trap closure. We observed two different scenarios. Either the two APs are elicited during the bending of the hair, or the first AP is generated during the bending, and the second while the sensory hair moves back into its original position. Below a minimal angular velocity of 0.03 rad/s, we never observed an AP and the trap consequently stayed open.

Taken together, our results show that a increasing strain on either side of the hair where the sensory cells are located, is the determining factor for the generation of APs. We propose that the mechanosensitive channels in the plasmamembrane of the sensory cells are open as long as they are under constant strain. Since this effect is counteracted by a membrane’s natural tendency to relax (the global effect is seen in the sharp force decay in our sustained displacement experiment), an increasing strain on the sensory hair is required to sustain the channels’ opening. As a consequence, the mechanosensitive channels will not open if the strain rate is too low (very slow deflection or sustained displacement). In a natural situation, when the deflection is fast and the angular displacement large enough, i.e. when an insect touches the sensory hair, the RP threshold is reached and a single AP is generated. In the case of a slower deflection, the RP threshold is reached during the deformation, eliciting an AP while the membrane is still under strain, thus reaching a second RP threshold during the same deflection. A second AP and trap closure are the consequence.

We can only speculate about the relevance of the observed single touch snapping in nature. However, one can imagine that it could be an advantage for catching slower prey animals, such as insect larvae that otherwise may not touch a hair twice within a 30 s time span.

## Acknowledgements

We would like to thank Karl Huwiler and Christian Frey for taking care of the Venus flytraps and providing us with healthy plants. This work was funded by ETH Zurich and the University of Zurich, and an interdisciplinary grant from the Swiss National Science Foundation (Grant Number CR22I2_166110) to U.G., B.J.N, and H.J.H.

## Author contributions statement

U.G. initiated the project. U.G., B.J.N., and H.J.H. conceptualized and supervised the project, and raised funding. J.T.B., E.S., N.F.L.. and H.V. designed, conducted, and analyzed the experiments. J.T.B. wrote the control and readout software. E.S., F.K.W., and F.R. developed the computational model. I.B. provided know-how and suggestions. J.T.B., E.S., N.F.L., and H.V. wrote the manuscript with help from U.G. All authors reviewed and commented on the manuscript.

