## Supplemental Figures for "The mechanical basis for snapping of the Venus flytrap, Darwin’s ‘most wonderful plant in the world’"

### Supplementary material

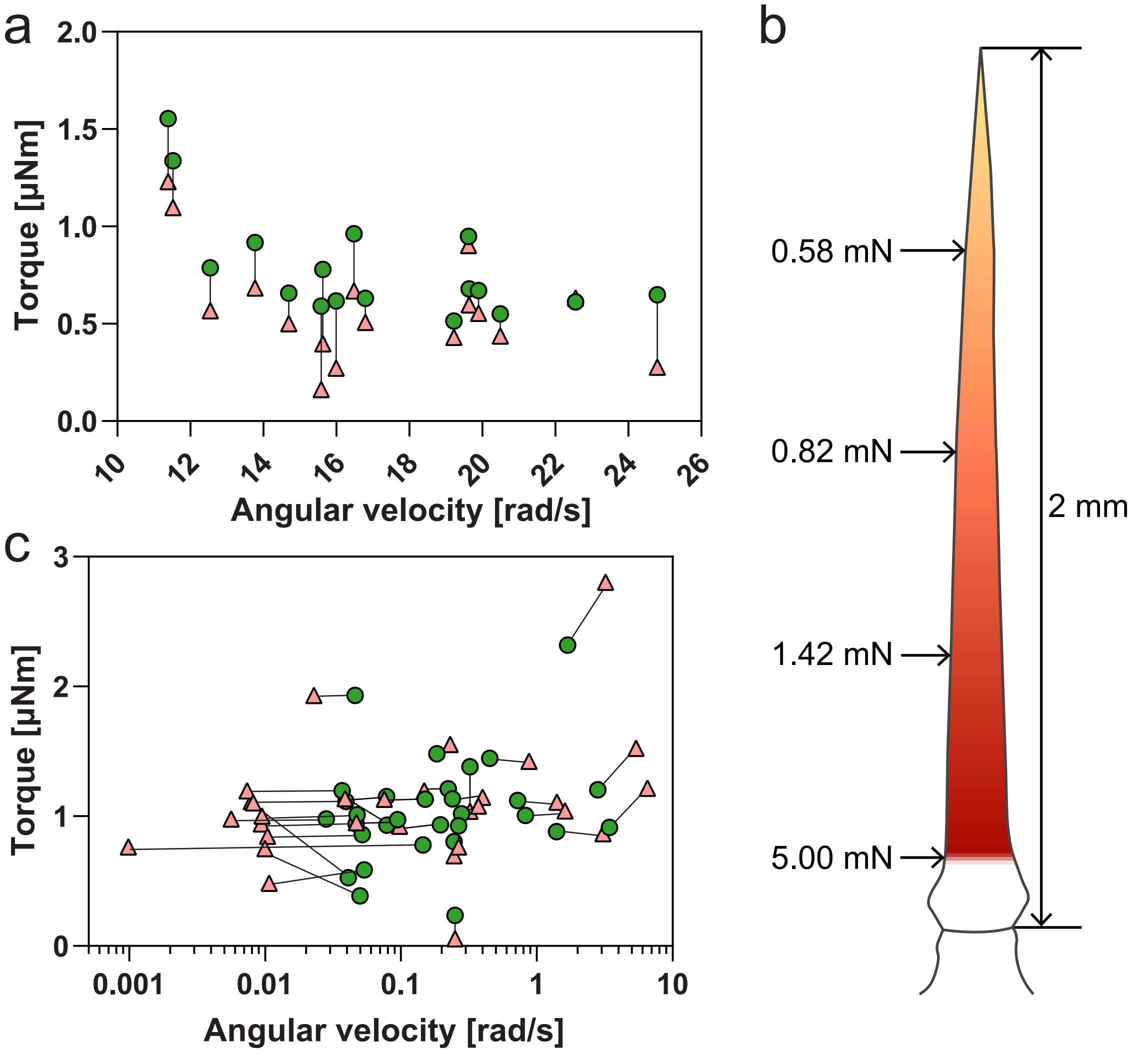


**Supplementary Figure 1.** (a) Torque vs. angular velocity plot for the double deflection experiments shown in figure 3b. (b) (e) Visualization of the needed force to sufficiently deflect the hair for double deflection triggering. (c) Torque vs. angular velocity plot for the single deflection experiments shown in figure 5b.

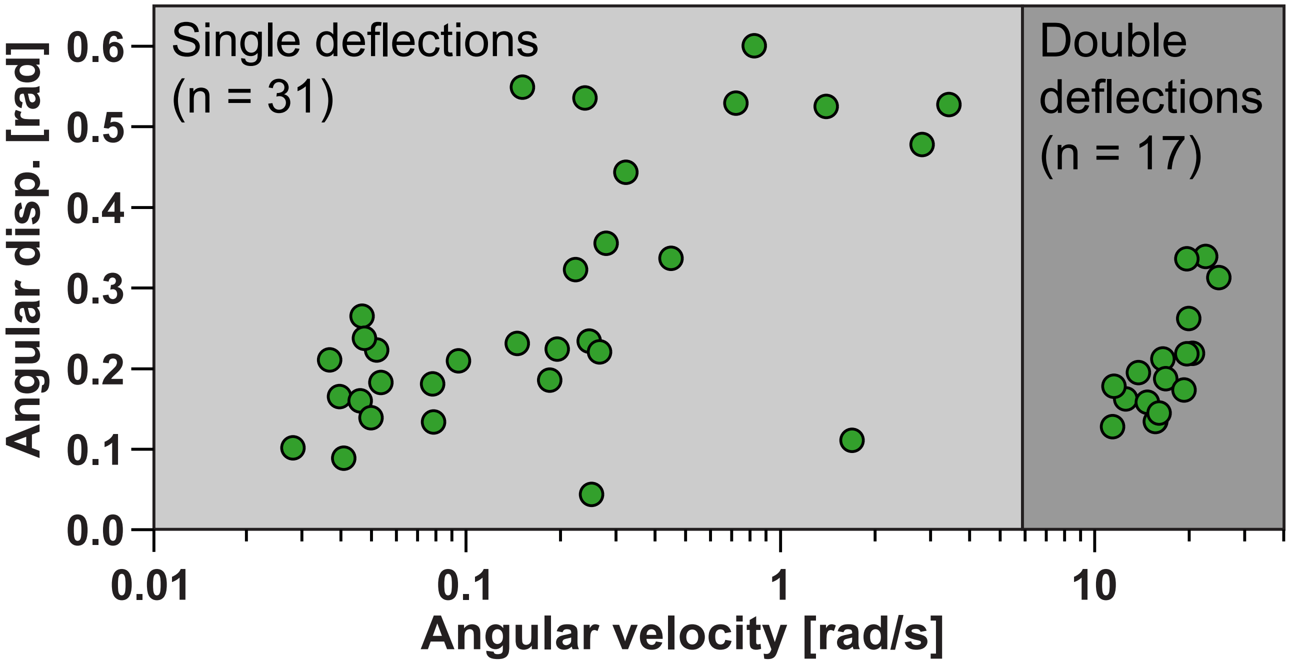


**Supplementary Figure 2.** All the mechanical stimuli (single and double deflections) that resulted in trap closure.
